# Bioinformatically predicted emulsifying peptides and potato protein hydrolysate improves the oxidative stability of microencapsulated fish oil

**DOI:** 10.1101/2022.11.18.517034

**Authors:** Mads Bjørlie, Betül Yesiltas, Pedro J. García-Moreno, F. Javier Espejo-Carpio, Nor E. Rahmani-Manglano, Emilia M. Guadix, Ali Jafarpour, Egon B. Hansen, Paolo Marcatili, Michael T. Overgaard, Simon Gregersen Echers, Charlotte Jacobsen

## Abstract

The aim of this study was to investigate the potential of potato proteins and peptides as emulsifiers in the microencapsulation of fish oil by spray-drying. Microcapsules were produced using a potato protein extract, and fractions enriched in patatin and protease inhibitors. Furthermore, bioinformatically predicted emulsifier peptides from abundant potato proteins and a hydrolysate, obtained through targeted proteolysis of the extract, were investigated. During 28 days of storage at 25°C, peptides and hydrolysate exhibited better emulsifying properties and higher encapsulation efficiencies compared to native proteins and sodium caseinate. Significant differences (p < 0.05) were observed in the peroxide value (PV) and secondary volatile oxidation products between the microcapsules produced with peptides and native proteins. Microcapsules produced with peptides and hydrolysate showed the highest oxidative stability, not exceeding a PV of 10 meq/kg oil, and with concentrations of volatiles below the odor threshold in oil for five of the six studied compounds. These results show the emulsifying potential of potato peptides and hydrolysate for use in microencapsulation of hydrophobic bioactive ingredients such as fish oil.

## 1 INTRODUCTION

Long-chain polyunsaturated fatty acids (PUFA) such as eicosapentaenoic (EPA, C20:5n-3), docosahexaenoic (DHA, C22:6-n3) and docosapentaenoic acid (DPA, C22:5-n3) have several, well-documented health benefits. They may prevent certain types of cancer, inflammations, cardiovascular diseases as well as improving the development and function of the central nervous system (Calder, 2017; Trautwein, 2001). This emphasizes the importance of EPA and DHA in the diet. The European Food Safety Authority recommends a daily dose of 250 mg EPA plus DHA (EFSA, 2017). However, the average daily amount of n-3 PUFAs in a western diet is around 150 mg (Beindorff & Zuidam, 2010). Fish oil is one of the richest sources for n-3 PUFAs and one of the ways to increase the amount of n-3 PUFAs in the diet could be to utilize fish oil supplements or foods enriched with fish oil (Kolanowski et al., 2006). One of the major obstacles of using fish oil in food enrichment is the polyunsaturated nature of PUFAs. The fatty acids are very prone to oxidation due to bis-allylic hydrogens in the chain, which leads to rancidity thus decreasing the nutritional and organoleptic properties. This limits the possibilities of incorporation of n-3 PUFAs into complex food matrices. Using microencapsulated omega-3 oils could potentially be a solution. For instance, (Jacobsen, 2015) reported that the peroxide value (PV) of energy bars enriched with microencapsulated fish oil remained stable compared to those enriched with neat fish oil in which the PV increased dramatically over the first 8 weeks of storage.

Microencapsulation is a technique that is already widely used in food, chemical, and agricultural industries. It is the process of enclosing small particles within a layer of coating or within a matrix. Microencapsulation is done to improve the shelf-life of the encapsulated material by limiting exposure to degrading environmental factors such as water, oxygen, heat, and light and to improve incorporation into complex food matrices that might contain pro-oxidants such as trace metals (Jacobsen, 2015). Microencapsulation can be done in multiple ways and the most common technique used in the food industry is spray-drying (Gibbs et al., 1999). Encapsulation is carried out by spray drying an emulsion that contains the core material (e.g. fish oil) dispersed in an aqueous phase with the dissolved encapsulating agent. The emulsion must be stable with small droplet size to ensure that the oil droplets are sufficiently covered by the interfacial film, which should result in a high encapsulation efficiency (Gaonkar et al., 2014; Soottitantawat et al., 2003). Carbohydrates are typically used as shell materials. However, carbohydrates often lack the interfacial properties of emulsifying agents and cannot create stable emulsions by themselves. Therefore, surface-active/amphiphilic compounds (e.g., proteins and peptides) with emulsifying activity are needed (Hogan et al., 2001a, 2001b).

Potato protein is produced as a side stream in potato starch production. It is increasingly gaining attention as it functions as both a protein source for human consumption as well as a unique food ingredient (Schmidt et al., 2018). Potato protein is extracted from potato fruit juice, which remains after starch and fibers have been extracted. Extraction was traditionally done using acid/heat treatment, but more gentle technologies are gaining attention to avoid denaturing the proteins. Despite the modest protein content (1-2%), the potato starch industry potentially produces 240,000 tons of high-quality protein per year, which can be used as ingredients in the food industry. Potato proteins have several functional properties of interest such as stabilizing foams and emulsions, gelling, and acting as antioxidants (Alting et al., 2011; García-Moreno, Jacobsen, et al., 2020; Schmidt et al., 2018). Potato protein composition is typically divided into three groups: patatins (up to 40%), protease inhibitors (up to 50%), and other proteins such as oxidative enzymes and enzymes involved in starch synthesis (Schmidt et al., 2018). Through enzymatic hydrolysis, the functional properties (e.g., antioxidant and emulsifying) can be enhanced as the functional groups on the proteins become exposed and their size is reduced (Cheng et al., 2010). Previous studies have shown that potato peptides and proteins can be utilized as emulsifiers with good results (García-Moreno, Gregersen, et al., 2020; García-Moreno, Jacobsen, et al., 2020; Schmidt et al., 2018; Yesiltas et al., 2021). García-Moreno, Gregersen, et al., 2020 used bioinformatics to identify specific peptide sequences in patatin and protease inhibitors with potential emulsifying activity based on their amphiphilicity in a given conformation. Through *in vitro* functional investigations, the authors were able to identify a range of novel peptides with strong emulsifying properties. These peptides were subsequently investigated to determine their secondary structure at the oil/water interface and obtain further insight on the properties of the interfacial layer (García-Moreno et al., 2021b). Recently, the structure/function relationship for these peptides were used as the basis for designing a targeted hydrolysis process, where application of trypsin was found to both release targeted peptides and improve the interfacial properties of the hydrolysate compared to hydrolysates produced using a range of other industrially relevant proteases (Gregersen Echers et al., 2022).

Considering the above, this study aimed to investigate the use of different potato proteins and peptides as emulsifiers in the microencapsulation of fish oil by spray drying. Six different proteins/peptides were examined; a food-grade potato protein extract (PPE) obtained from a non-denaturing extraction method, a targeted enzymatic hydrolysate of PPE using trypsin, a purified patatin fraction and protease inhibitor fraction from PPE, as well as to synthetic peptides derived from major potato proteins, γ-1 (GIKGIIPAIILEFLEGQLQEVDNNKDAR) and γ-38 (FCLKVGVVHQNGKRRLALVKDNP), both with validated emulsifying activity (García-Moreno et al., 2021b; Yesiltas et al., 2022). The physical characterization of the emulsions and capsules and the oxidative stability of the capsules during 28 days of storage were evaluated.

## 2 MATERIALS AND METHODS

### 2.1. Materials

Commercial cod liver oil was provided by Vesteraalens A/S (Sortland, Norway) and stored at –40ºC until use. Sodium caseinate (Cas) was provided by Arla Foods Ingredients amba (Viby J, Denmark). Food-grade potato protein extract (PPE) from non-denaturing extraction was provided by KMC amba (Brande, Denmark). Despite the high protein content in the potato protein extract we have chosen to use the term PPE rather than potato protein isolate (PPI) although the latter term is often used for extracts with high protein contents. The patatin and protease inhibitor content of the extract was 29-35% and 53-58% respectively (García-Moreno, Gregersen, et al., 2020). Patatin-rich (Pat) and protease inhibitor-rich (PI) fractions of the PPE were produced by Lihme Protein Solutions (Lyngby, Denmark) and obtained using gentle sequential precipitation through a proprietary pH-shift methodology. The synthetic peptides γ-1 (GIKGIIPAIILEFLEGQLQEVDNNKDAR) and γ-38 (FCLKVGVVHQNGKRRLALVKDNP) were purchased from ChinaPeptides Co., Ltd (Shanghai, China). The molecular weight was 3.094 kDa and 2.592 kDa for G1 and G38, respectively. The peptide purity (by HPLC, supplier certified) was 84.52% and 88.73% for G1 and G38, respectively. Glucose syrup (DE38, C*Dry 1934) was donated by Cargill Germany GmbH (Krefeld, Germany). An enzymatic hydrolysate of the potato protein extract (PPH) was prepared using pancreatic trypsin (Novozymes A/S, Bagsværd, Denmark) to a degree of hydrolysis of 5.4% (based on alpha-amino nitrogen using the ninhydrin method), corresponding to an average peptide chain length of 18.1 (Gregersen Echers et al., n.d.). All other chemicals and solvents used were of analytical grade.

### 2.2. Preparation of the emulsion

The seven different emulsions were produced using the protein-based emulsifying agents and the same encapsulating agent (glucose syrup). A 10mM imidazole/sodium acetate buffer at pH 7 was used. The emulsifying agent (0.2% w/w) was dissolved in the buffer after which the glucose syrup (28% w/w) was added to the buffer. The resulting aqueous phase was stirred overnight at 250 rpm. A pre-homogenization step was carried out for three min at 18,000 rpm using a T25 digital ULTRA-TURRAX (IKA) during which the oil (5% w/w) was added during the first minute. Homogenization was carried out in a high-pressure, 2-valve homogenizer (Panda 2K, Niro Soavi Deutschland, Lübeck, Germany) using a pressure profile of 450/75 bar and passing the emulsion through three times. The temperature of the emulsion was controlled using an ice-cooled water bath.

### 2.3. Spray-drying the emulsion

The emulsions were spray-dried in a pilot plant scale spray-drier (Mobile Minor, Niro A/S, Copenhagen, Denmark). The emulsions were dried using inlet/outlet temperatures of 190°C and 80°C respectively. The pneumatic air pressure, which controls the speed of the rotary atomizer, was set to four bar which results in rotational speed of 22,000 rpm. The emulsions were stirred in a magnetic stirrer at 250 rpm while being pumped to the spray-drier to avoid potential physical destabilization such as creaming.

### 2.4. Characterization of the emulsion

Oil droplet size distribution (ODSD) was measured using a Mastersizer 2000 (Malvern Instruments, Ltd., Worcestershire, UK). ODSD was measured for both the original emulsion as well as the reconstituted emulsion (obtained by dispersing the capsules in water) to investigate potential differences. Emulsions were reconstituted to the original water content of 66.8%; one g of powder was mixed with 2.01 g of water and vortexed briefly. The measurements were carried out as described by (García-Moreno et al., 2018). Solutions were diluted in recirculating water (3000 rpm) until they reached an obscuration of 12%. The refractive indices of sunflower oil (1.469) and water (1.330) were used as particle and dispersant, respectively. Measurements were made in triplicate and the 90^th^ percentile (D_90_), as well as the surface-weighted mean diameter (D_3,2_), and the volume-weighted mean diameter (D_4,3_) were reported.

### 2.5. Characterization of the spray-dried powder

#### 2.5.1. Capsule morphology

The morphology of the capsules was investigated using scanning electron microscopy (SEM) (FEI Inspect, Hillsboro, OR, USA). A thin layer of powder was placed on carbon tape and sputter-coated with gold, 8s, 40 mA using a Cressington 208HR Sputter Coater (Cressington Scientific Instruments, Watford, England). The capsule diameter size distribution was determined from the SEM images using the open-source image-processing program ImageJ (National Institutes of Health). One hundred capsules were selected at random for each of the samples.

#### 2.5.2. Encapsulation efficiency

Encapsulation efficiency (EE) was determined from the amount of easily extracted fat e.g. surface fat on the capsules. The fat was extracted according to (Danviriyakul et al., 2002) with some modifications. One g powder was added to 7.5 mL hexane and gently shaken for two min at 100 rpm. The samples were then centrifuged for 10 min at 3000g. Three mL of the supernatant were collected in a previously weighed Pyrex tube and the solvent was evaporated to dryness under a flow of nitrogen. The tubes were then placed in a 102°C oven for one hr and cooled down in a desiccator for 10 min. The Pyrex tubes were weighed again, and the oil concentration was adjusted to the initial volume of solvent. EE was calculated as shown below:

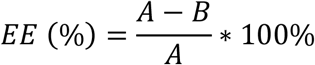

A is the theoretical amount of oil and B is the non-encapsulated oil. Measurements were carried out in triplicates.

#### 2.5.3. Moisture content

The moisture content was measured using an infrared balance (AD-471A, Tokyo, Japan). Approximately one g of powder was placed on the scale and heated for 90 minutes at 105°C until constant weight. Measurements were carried out in duplicates.

### 2.6. Oxidative stability

#### 2.6.1. Storage experiment and sampling

Immediately after production, the capsules were placed in a -80°C freezer. At the beginning of the storage experiment, the capsules were divided into 30 mL brown bottles (approximately 10 g in each bottle), sealed with plastic caps and placed in a 25°C oven. One bottle per sample per week for PV/tocopherols and volatiles respectively. Once a week the bottles were removed from the oven, purged with nitrogen and placed in a sealed plastic bag in a -40°C freezer until analysis.

#### 2.6.2. Determination of oil content and peroxide value

The oil content of the capsules was determined using the method described by (Bligh & Dyer, 1959) with some modifications. One g of capsules was weighed in a centrifuge tube and 15 mL distilled water was added. Thirty mL methanol and 15 mL chloroform were added to the glass followed by mixing for 20 s using an Ultra-Turrax mixer at 6500 rpm. Fifteen mL of chloroform was added followed by mixing for 20 s. After mixing, 15 mL of distilled water was added, and the tube was gently shaken. The tubes were centrifuged in a Sigma 4-16KS centrifuge (Sigma Zentrifugen, Osterode am Harz, Germany) at 2800 rpm and 18°C for 10 min. Following the centrifugation, the methanol/water phase was removed using vacuum suction. The remaining extract was filtered into a Pyrex bottle using phase separation paper. Six-eight g extract was weighed in a tared and previously weighed 25 mL beaker and placed in a fume hood for evaporation overnight. Any remaining water was removed in a 105°C oven for one h followed by cooling in a desiccator for 20 min. Two extracts were made from each sample. Peroxide value was determined on the lipid extracts using the colorimetric ferric-thiocyanate method at 500 nm as described by (Shantha & Decker, 1994). Results were expressed as milliequivalents of peroxides per kg of oil.

#### 2.6.3. Determination of tocopherol content

Tocopherol content was determined using high-performance liquid chromatography (HPLC) (Agilent 1100 Series; Column: Waters Spherisorb 3 μm Silica; 4.6 × 150 mm) using the same lipid extracts as the PV analysis. Two g of extract was weighed and evaporated under a flow of nitrogen. The oil was re-dissolved in one mL heptane and transferred to the injection bottles and sealed. HPLC analysis was carried out as described in AOCS Official Method Ce 8-89. Mobile phase was heptane:2-propanol in the ratio 100:0.4 (v/v). Injection volume was 20 μL and flowrate was 1.0 mL/min. Detection was done using a fluorescence-detector with excitation at 290 nm and emission at 330 nm. Measurements were carried out in duplicate.

#### 2.6.4. Determination of volatile secondary oxidation products

The volatile secondary oxidation products were determined by gas chromatography-mass spectrometry (GC-MS), as described by García-Moreno et al. (2016) with some modifications. One g of capsules was weighed in a purge bottle together with 30 mg of internal standard (4-methyl-1-pentanol, 30 μg/g water) and mixed with 10 mL of distilled water. The bottle was heated for 30 minutes in a 45°C water bath while purging with nitrogen (150 mL/min). The volatile secondary oxidation products were trapped on Tenax GR tubes. The volatiles were desorbed again by heating (200°C) in an Automatic Thermal Desorber (ATD-400, Perkin Elmer, Norwalk, CN), cryofocused on a cold trap (-30 °C), released again (220 °C), and led to a gas chromatograph (HP 589+IIA, Hewlett Packard, Palo Alto, CA, USA; Column: DB-1701, 30 m × 0.25 mm × 1.0 μm). The individual compounds were analyzed by mass spectrometry (HP 5972 mass-selective detector, Agilent Technologies, USA; electron ionization mode, collisional energy = 70 eV; MS1 scan mode (30-250 m/z)). The individual compounds were identified by MS-library searches (Wiley 138 K, John Wiley and Sons, Hewlett-Packard) and quantified using calibration curves made using external standards. These standards were 2-ethyl-furan, 1-penten-3-one, pentanal, 1-penten-3-ol, 2,3-pentanedione, (*E*)-2-pentenal, 1-pentanol, hexanal, (*E*)-2-hexenal, heptanal, (*Z*)-4-heptenal, benzaldehyde, octanal and (E,E)-2,4-heptadienal. The standards were dissolved in 96% ethanol and diluted to concentrations of approximately 2.5, 5, 10, 50, 100, 200, 500, and 1000 μg/mL. One μL of each solution was injected directly onto a Tenax tube. Results were expressed as ng/g oil and measurements were made in triplicate.

### 2.7 Statistical analysis

Data analysis was done using the Statgraphics Centurion 18 software (Statistical Graphics Corp., Rockville, MD, USA). Data were expressed as mean ± standard deviation. Firstly, multiple sample comparison analysis was performed to identify significant differences between the samples. Secondly, mean values were compared using Tukey’s test. Differences were considered significant at p < 0.05.

## 3 RESULTS AND DISCUSSION

To investigate the applicability of potato protein and derivates for the production of spray dried emulsions, we investigated a number of different protein-based substrates as emulsifiers. In addition to a native, food-grade protein isolate, we investigated two fractions enriched for patatin and protease inhibitors, respectively. Additionally, we included a potato protein hydrolysate obtained by proteolysis using trypsin, showing superior emulsifying properties from a benchmark study against hydrolysates produced using a range of industrial proteases (Gregersen Echers et al., 2022). Furthermore, we included two synthetic peptides, previously shown to have strong in vitro emulsifying properties (García-Moreno et al., 2021b; Yesiltas et al., 2022). γ-1 (GIKGIIPAIILEFLEGQLQEVDNNKDAR) is found in patatin, has a negative net charge at pH 7 and did not show in vitro antioxidant activity (DPPH radical scavenging, Fe^2+^ chelating ability and Fe^3+^ reducing power). Despite the lack of antioxidant activity, γ-1 had the best emulsifying activity and was selected for this study because of this. γ-38 (FCLKVGVVHQNGKRRLALVKDNP) is found in Kunitz type-B serine protease inhibitors, has a positive net charge at pH 7 and showed in vitro antioxidant activity (DPPH radical scavenging), and displayed excellent emulsifying properties. γ-38 was included to represent a different type of peptide emulsifier based on physicochemical and antioxidant properties.

### 3.1 Emulsion preparation and spray drying

PPH, G1, G38 and Cas produced emulsions that were physically stable from production until spray drying. The emulsions stabilized with PPE as well as Pat-, and PI-rich fractions (**Figure S2)** separated shortly after pre-homogenization and again after homogenization. Using magnetic stirring, it was possible to keep the fish oil sufficiently dispersed while pumping the emulsions to the spray-drier. Spray-dried fish oil-loaded microcapsules were obtained for all seven samples. The color of the powder varied from white to greyish depending on the color of the peptides or proteins. Synthetic peptides resulted in white powder while the PPH resulted in a darker greyish powder. Powder stuck to the side of the spray drier, which could be caused by oil on the surface of the capsules. The average oil content of the capsules, as determined by the Bligh & Dyer method, was 14.8 ± 2.3 %. The theoretical oil content of the capsules was 15.1 dm wt% indicating that almost all the oil was encapsulated.

### 3.2. Oil droplet size distribution

Emulsion droplet size has a great impact on the potential to microencapsulate the oil using spray drying. (Soottitantawat et al., 2003) reported that smaller droplet sizes resulted in less surface oil, better flavor retention, and smaller capsules when encapsulating volatile compounds. Oil droplet size depends on the surface-active properties of the emulsifier and thus gives an indication of the potential of the protein or peptide as an emulsifier. For microencapsulation through spray-drying, the 90^th^ percentile (D_90_) of the capsule diameter should be below 2 μm (Tamm et al., 2015). The Oil droplet size distribution (ODSD) for both the parent emulsion and the reconstituted emulsions are shown in **Table 1**. Sodium caseinate has been used as a stabilizer for microencapsulation via spray-drying in previous studies with good results (Hogan et al., 2001a, 2001b; Ixtaina et al., 2015). In the present study, it was found that Cas exhibited sufficient emulsifying activity (D_90_ < 2 μm), even at the low concentration of 0.2%. G1, G38, and PPH showed emulsifying activity similar to Cas with D_90_ and D_4,3_ values which were not significantly different from Cas. Additionally, G1, G38, and PPH all achieved D_90_ values below 2 μm indicating sufficient stability for microencapsulation. Pat, PI and PPE all showed lower emulsifying activity with D_90_ droplet sizes well above the 2 μm limit. (Tamm et al., 2015) achieved D_90_ values between 1.40 and 1.64 μm when encapsulating fish oil using various hydrolysates of beta-lactoglobulin as stabilizer and glucose syrup as the encapsulating agent. The emulsion used in their study contained 29.5 wt% glucose syrup, 14.5 wt% oil and 2.2 wt% emulsifier resulting in an oil/emulsifier ratio of 6.6 compared to 25 in the present study indicating that G1, G38, and PPH are more efficient at emulsifying than beta-lactoglobulin hydrolysates. Cheng et al., 2010 investigated the potential of Alcalase hydrolyzed potato protein as an emulsifier and antioxidant in soybean oil-in-water emulsions. The emulsions contained 10 wt% oil and 0.25 to 2 wt% emulsifier resulting in an oil/emulsifier ratio between 40 and 5. They found that the hydrolysate was not able to stabilize the emulsions by itself and therefore had to be used in conjunction with Tween 20. In comparison, both G1, G38 and PPH were able to stabilize emulsions without additional emulsifiers, although at an oil/emulsifier ratio of 25. This clearly highlights the importance of the applied protease for hydrolysis, and that targeted hydrolysis, here using trypsin, <u>can</u> facilitates production of hydrolysates with superior emulsifying properties compared to conventional trial-and-error hydrolysates.

Reconstituting the emulsions showed a significant change in D_4,3_ for all samples except for G38 and PPE. D_4,3_ is sensitive to larger oil droplets and an increase thus suggests coalescence of the droplets. Furthermore, a difference in droplet size between parent and reconstituted emulsions reflects instability during atomization (Tamm et al., 2015). Indeed, an increase in both D_90_ and D_4,3_ was observed for the three emulsions that broke after homogenization. However, the increase in both D_90_ and D_4,3_ was unexpected for Cas and PPH as these emulsifiers provided physically stable emulsions. Only a very small increase in D_4,3_ was observed for G1 indicating excellent emulsifying activity with minimal droplet coalescence. During spray-drying, the surface-active peptides remain at the interface of the oil droplets between the oil and the shell material (Tamm et al., 2015). Non-encapsulated oil is therefore not stabilized with emulsifiers and can coalesce easier which could lead to droplet aggregation when reconstituting the emulsion. The differences observed in emulsifying activity for the different potato protein/peptide samples can be attributed to several different factors. Emulsifiers have two primary functions: 1) to decrease the interfacial tension between the oil and water phases to enable disruption of the droplets during homogenization and 2) to form a protective coating around the droplets to prevent coalescing (McClements, 2004). Additionally, there must be enough emulsifier to cover the droplet surfaces and the emulsifiers must be allowed enough time to move from the bulk phase to the droplet surface. Proteins that are used as emulsifiers must be amphiphilic in order to adsorb at the oil-water-interface interface. This amphiphilicity is a result of hydrophobic regions or amino acids that can be buried inside of larger proteins.

**Table 1.**
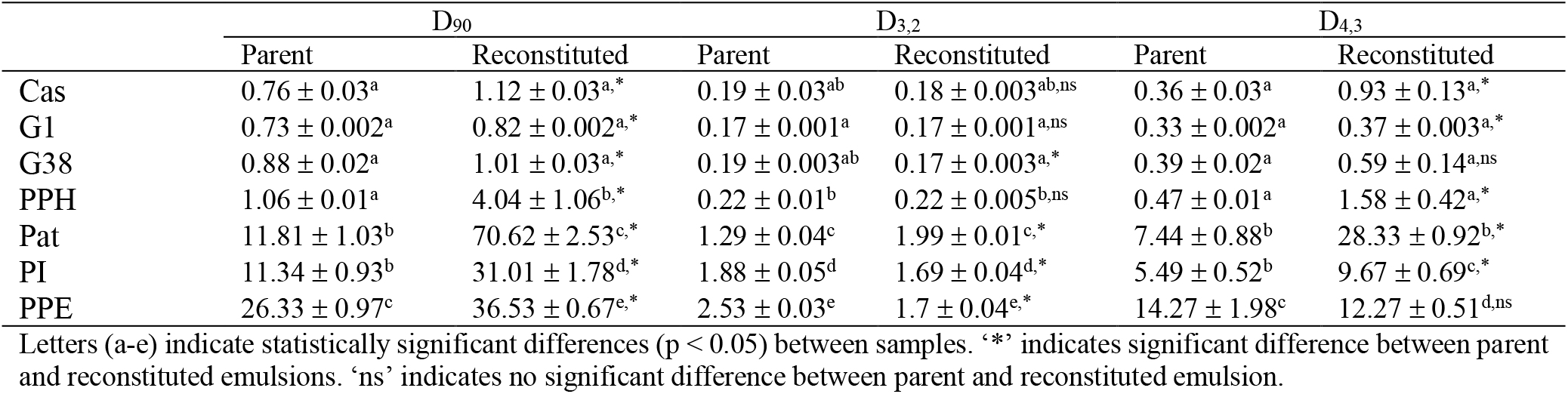
Oil Droplet Size Distribution for parent and reconstituted emulsions.

Through hydrolysis, these hydrophobic regions or amino acids can be exposed which increases the net surface amphiphilicity of the resulting peptides (McClements, 2004). G1 and G38 both have several of the hydrophobic groups in their sequence and has been shown to exhibit their emulsifying activity through the formation of an amphiphilic α-helix and β-strand, respectively (García-Moreno et al., 2021a). The increased emulsifying activity of the peptides and hydrolysate compared to Pat, PI and PPE could therefore possibly be attributed to a larger relative exposure of hydrophobic groups in the peptides or formation of more energetically favorable amphiphilic secondary and tertiary structures. Another possible explanation for the lower emulsifying activity of PI, Pat, and PPE could be insufficient adsorption time for the proteins to diffuse from the water to the oil. Larger proteins (40-45 kDa for Pat and 10-24 kDa for PI (García-Moreno, Gregersen, et al., 2020)) take longer time to diffuse to the interface than peptides (even if both are amphiphilic) due to differences in size (Drusch et al., 2007). This can result in a high concentration of proteins in the aqueous phase. In a previous study, (Schmidt et al., 2018) reported the emulsifying activity of both potato protein powder, purified patatin, and PI, and fractions of patatin and PI separated based on hydrophobicity when emulsifying sunflower oil. They found that the emulsifying activity index of potato protein powder was not significantly different from purified patatin, but higher than that of purified PI. The high hydrophobicity fractions of patatin and PI showed improved emulsifying activity while the low hydrophobicity fraction of PI had decreased emulsifying activity. This is in line with the results in the present study indicating improved emulsifying activity when exposing hydrophobic amino acids in the peptides. However, this does not explain the inability of the potato proteins to form stable emulsions in the present study. This could be attributed to a combination of the size of the proteins and the increase in viscosity of the aqueous phase when adding the glucose syrup as it further decreases the diffusivity of the proteins. Finally, the presence of carbohydrates can also affect homogenization negatively (McClements, 2004), however, the same shell material was used for all of the samples and therefore the potential effect would be due to an interaction between glucose syrup and Pat, PI or PPE.

### 3.3. Capsule morphology

**Error! Reference source not found.** shows the scanning electron micrographs of the seven different fish oil-loaded microcapsules. Spherical capsules were obtained for all seven emulsifiers. The capsules showed little to no agglomeration which can be caused by oil on the surface of the capsules causing them to stick together (Hogan et al., 2001b). Cas, G1, and G38 produced capsules with the smoothest surfaces compared to Pat, PI, and PPE. Capsules produced with PPH generally had smooth surfaces with some of the capsules displaying a rougher, textured surface. It is important to note that PPH is a mixture of peptides containing both emulsifying peptides and peptides with potential antagonistic or interfering effects. (Hermund et al., 2019) also obtained capsules with rough surfaces and some agglomerates when using glucose syrup as the shell material compared to dextran. PI (**Figure 1F**) showed pores inside the shell material which could have been the result of oxygen inclusion during homogenization and trapping of air bubbles during spray-drying (Drusch et al., 2009). Some of the capsules, especially G38, Pat and PI, displayed black spots on the surface which may indicate oil droplets on the surface (Drusch & Berg, 2008). The seven types of microcapsules presented a broad range of sizes varying from two to 40 μm. The mean diameters of the capsules produced using Pat, PI, and PPE as emulsifiers were significantly larger than the other four types. Capsule mean diameter ranged from 7.2 to 15.4 μm for PPH and Pat respectively. For Cas, G1, and G38 approximately 70% of the capsules were below 10 μm while for PPH it was approximately 85%. Conversely, 80% of the capsules were above 10 μm for the Pat, PI and PPE capsules. The spray-dried capsules obtained in this study showed size distributions comparable to, or smaller than, previously reported results. (Drusch, 2007) reported mean diameters from 14.2 to 18.1 μm when using a mixture of sugar beet pectin and glucose syrup as encapsulating agents and oil loads of 50 and 20 dm wt% while (Carneiro et al., 2013) reported mean diameters between 17.9 and 23 μm in microencapsulation of flaxseed oil using various shell materials and an oil load of 20 dm wt%. As pointed out by (García-Moreno et al., 2018) capsules with reduced size are preferable, when the goal is to incorporate them into food matrices. Smaller capsules can be easier to disperse and have a reduced effect on the texture of the product compared to larger capsules. Additionally, the smaller the diameter of the capsules, the larger the surface area to volume ratio. This presents a dilemma as the large surface can enhance the release of bioactive compounds, but at the same time, a large surface-to-volume ratio increases the contact surface between lipids and prooxidants which can have a negative impact on the oxidative stability (García-Moreno, Jacobsen, et al., 2020).

**Figure 1.**
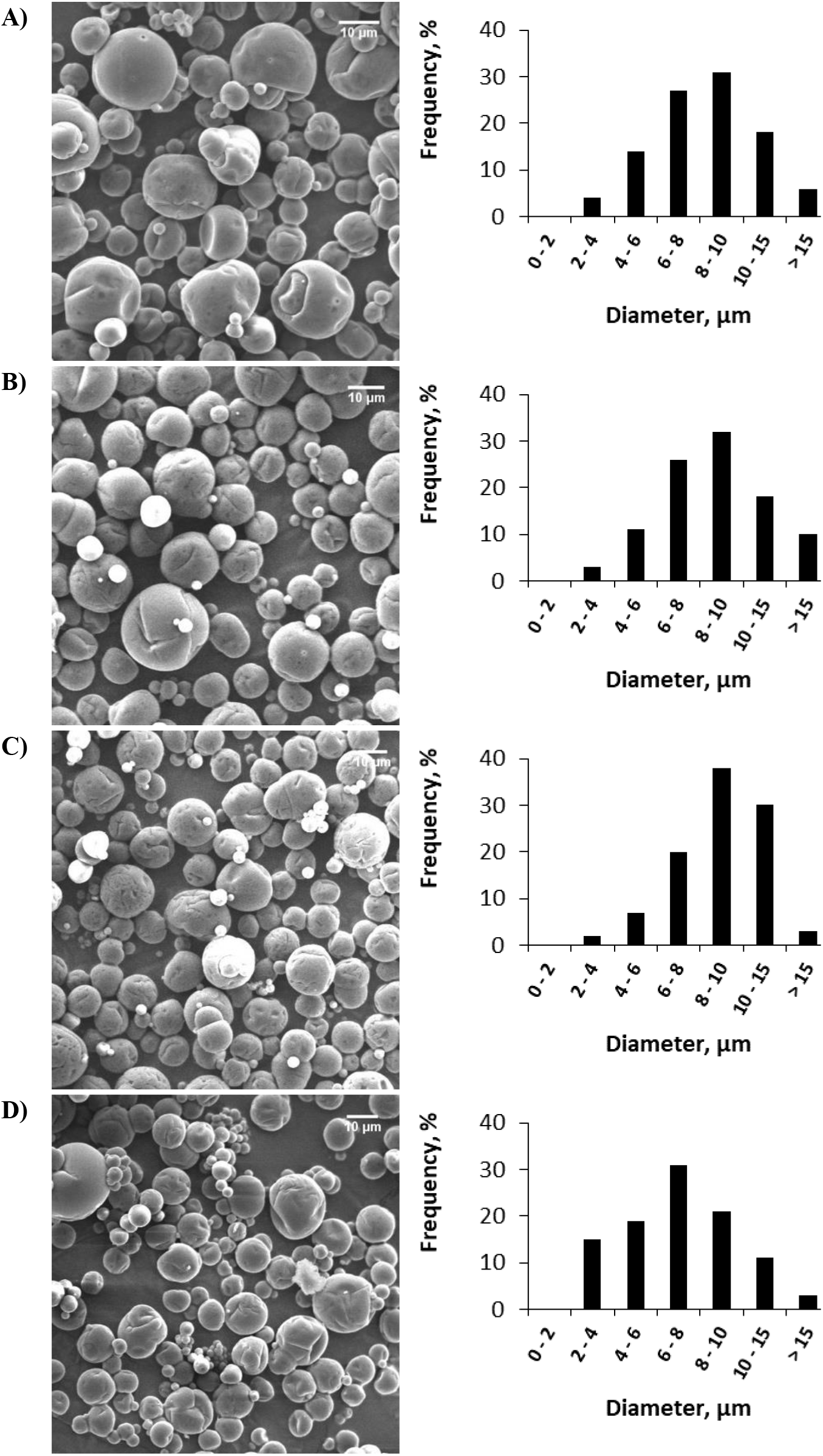
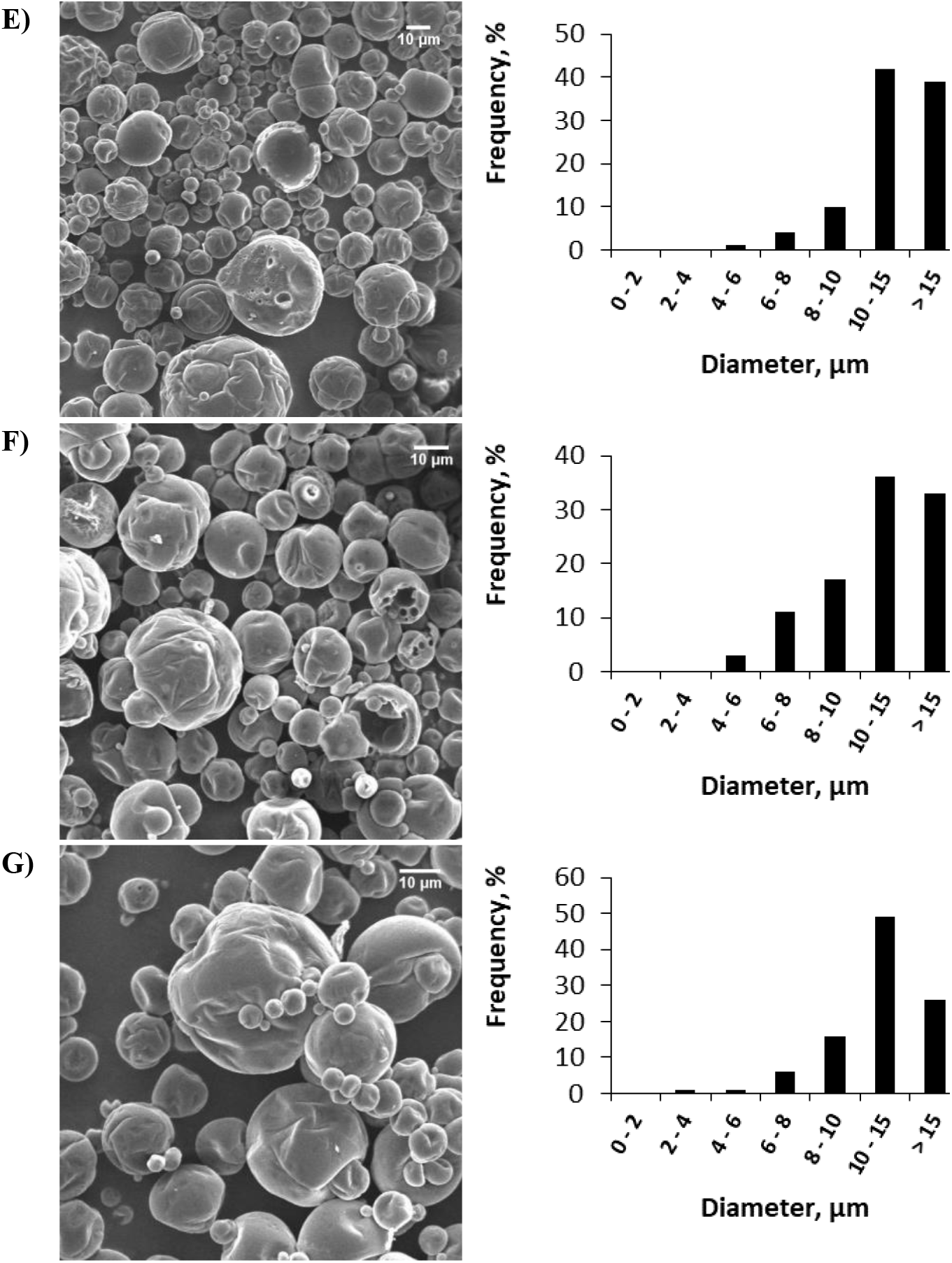
SEM images and size distributions of the seven different fish oil-loaded spray-dried microcapsules. A) Cas, B) G1, C) G38, D) PPH, E) Pat, F) PI, G) PPE

### 3.4 Encapsulation efficiency and moisture content

Non-encapsulated oil or ‘surface oil’ is a critical quality parameter in microencapsulation of oils by spray-drying. Oil on the surface of the capsules adversely affects the flowability and wettability of the spray-dried powders (Vega & Roos, 2006). Additionally, and even more critical, non-encapsulated oil is not protected by the encapsulating agent and is more prone to oxidation. Unprotected polyunsaturated fatty acids, such as those found in fish oil, may strongly affect the product acceptability through the formation of volatile secondary oxidation products, which can result in undesired odors and off-flavor (Drusch & Berg, 2008). Encapsulation efficiency (EE) is a measure of how much of the total oil that has been encapsulated inside the glassy matrix. The analytical method used to determine the amount of surface oil can influence the results (Drusch & Berg, 2008) and varies from study to study. (Danviriyakul et al., 2002) used a 2 min vortex followed by 20 min centrifugation while (Hermund et al., 2019) used gentle shaking followed by gravitational filtration. Centrifugation, as done in the present study, could potentially disrupt the capsule structure, which would allow the solvent to reach the encapsulated oil resulting in an overestimation of the EE. The results should therefore be interpreted carefully. **Table 2** shows the encapsulation efficiencies for the seven types of fish oil-loaded microcapsules. Efficiencies ranged from 62.2 ± 6.8 to 93.1 ± 0.7% for PPE and G1, respectively. Significantly higher efficiencies were obtained for G1 and G38 compared to Pat, PI, and PPE. G1 also displayed a significantly higher efficiency compared to Cas. Interestingly, PPH was not significantly different from the G1, G38 or Cas. The efficiencies obtained in the present study compare favorably with efficiencies previously reported. (Hermund et al., 2019) reported encapsulation efficiencies from 83.2 to 90.4%, while (García-Moreno et al., 2018) reported efficiencies 78.1 and 91.7%. Both studies concerned electrospraying which is another technique used in microencapsulation. (Tamm et al., 2015) reported a mean EE of 99 ± 0.5% for microencapsulation of fish oil by spray-drying using whey protein as stabilizers. Their study used an oil load of 14.5% and 2.2% emulsifier (6.6:1) in emulsion compared to the 5% oil load and 0.2% emulsifier (25:1) used in the present study and can thus not be directly compared for evaluation of efficiency. Furthermore, (Hogan et al., 2001a) showed that the ratio between emulsifier and the encapsulating agent had a significant impact on the encapsulation efficiency obtaining efficiencies between 62 and 92% for 1:69 and 1:19 respectively. The ratio used in the present study is 1:140. These arguments suggests that higher efficiencies might be attainable if using a larger amount of emulsifier. Nevertheless, our results show that peptide-based emulsifiers from potato efficiently encapsulate fish oil – even at low dosage. The moisture content of the fish oil-loaded capsules is shown in **Table 2**. The moisture contents varied between 3.4 and 4.4%, indicating that the drying was properly carried out. The moisture content was not affected by the different emulsifiers used.

**Table 2.** Moisture content and encapsulation efficiency

|  | Moisture content, % | Encapsulation efficiency, % |
| --- | --- | --- |
| Cas | 4.4 ± 0.5<sup>a</sup> | 80.7 ± 1.0<sup>bc</sup> |
| G1 | 3.9 ± 0.1<sup>a</sup> | 93.1 ± 0.7<sup>a</sup> |
| G38 | 4.4 ± 0.4<sup>a</sup> | 90.5 ± 0.3<sup>ab</sup> |
| PPH | 3.6 ± 0.5<sup>a</sup> | 84.6 ± 0.5<sup>abc</sup> |
| Pat | 3.4 ± 0.6<sup>a</sup> | 75.0 ± 0.8<sup>cd</sup> |
| PI | 3.8 ± 0.3<sup>a</sup> | 64.6 ± 7.3<sup>de</sup> |
| PPE | 4.4 ± 0.6<sup>a</sup> | 62.2 ± 6.8<sup>e</sup> |

### 3.5. Oxidative stability

The oxidative stability of the fish oil-loaded capsules was evaluated by measuring the formation of primary and secondary volatile oxidation products as well as the α-, δ- and γ-tocopherol content during 28 days of storage at 25°C.

#### 3.5.1. Peroxide value

**Figure 2A** shows the peroxide values (PV) and tocopherol contents of the seven different fish oil-loaded capsules during storage. The PV after production (day 0) ranged from 1.1 ± 0.2 to 2.8 ± 0.1 meq/kg oil. All samples except for G38 and Pat showed significantly higher PV than that of the fresh fish oil (0.3 ± 0.01 meq/kg oil). Previous studies have attributed the change in PV during microencapsulation to oxygen inclusion and increase in specific surface area during emulsion preparation (Serfert et al., 2009) and exposure of the non-encapsulated oil to atmospheric air during the spray-drying process (Drusch et al., 2006). After 28 days of storage, the PV had increased significantly for all seven samples with final PV varying from 6.0 ± 0.2 to 188 ± 28 meq/kg oil. It must be noted that the method is valid for oils with a maximum PV of 40 meq/kg oil and therefore the values above 40 should be interpreted carefully. The type of emulsifier had a significant impact on the oxidative stability. Emulsions stabilized with Pat, PI or PPE showed no lag phase compared to the other emulsifiers (see **Table S10** in Supplementary Materials). The first sampling point, at which the PV is significantly different from that of day zero, determines the end of the lag phase. The lag phase of Cas and PPH lasted one week while G38 exhibited a lag phase of three weeks. The measurement at day 14 for G1 seems high compared to day 21 and day 28. It is not unusual to see an increase followed by a decrease during storage. However, the decrease of PV is related to the decomposition of hydroperoxides and the formation of volatiles. Given that there were no sudden increases in concentrations of volatiles at day 14 for G1 (**Figure 2A**) this increase can be disregarded and the lag phase is, therefore, three weeks. Both G1 and G38 had longer lag phases compared to Cas. Overall, G1, G38, and PPH showed little increase in PV during the 28 days of storage. Indeed, no significant differences in final PV were observed between G1, G38, Cas and PPH. According to the European Pharmacopoeia, the PV of fish oil should not exceed 10 (Hu & Jacobsen, 2016). G1, G38, and PPH were the only samples with PV below 10 after 28 days of storage, showing that microcapsules produced with peptides and the hydrolysate all displayed significantly higher oxidative stability than their parent proteins based on the PV. This is in line with the ODSD and the EE results where the peptides and hydrolysate performed better than the proteins. (Hermund et al., 2019) reported PV values from 20 ± 4.5 to 35 ± 0.5 meq/kg oil after 21 days of storage for fish oil-loaded electrosprayed capsules using glucose syrup or dextran as wall materials and a combination of seaweed extracts and commercial antioxidants. (García-Moreno et al., 2018) obtained PV values of approximately 20 meq/kg oil after 21 days of storage for fish oil-loaded electrosprayed capsules using dextran and glucose syrup as wall materials and whey protein as an emulsifier. (Drusch & Berg, 2008) reported PV values of approximately 120 meq/kg oil after 28 days of storage for fish oil-loaded spray-dried capsules (30 wt%) using a mixture of n-OSA starch and glucose syrup as wall materials. The PV obtained in this study for PPH, G1 and G38 were lower than, or comparable to, values reported in previous studies indicating potential as emulsifiers used in microencapsulation of fish oil.

**Figure 2.**
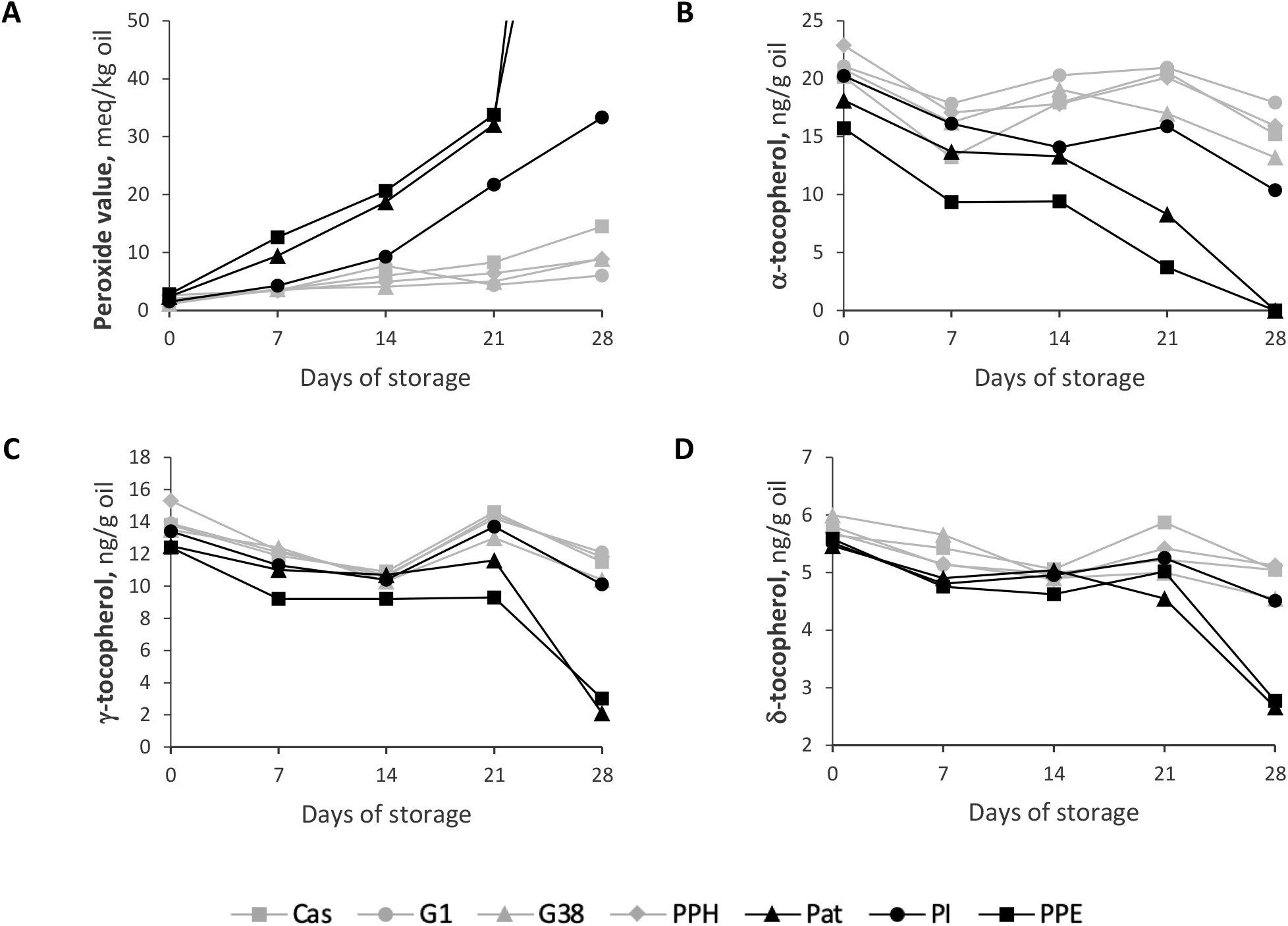
Peroxide value and tocopherols consumption during 28 days of storage for fish-oil, which has been microencapsulated using sodium caseinate, peptides or proteins as emulsifying agents. A) Peroxide value, B) α-tocopherol, C) γ-tocopherol, D) δ-tocopherol. Peroxide value at day 28 of Pat and PPE reached 188 and 127 meq/kg oil, respectively.

#### 3.5.2. Tocopherol content

Tocopherols are important lipid-soluble, natural antioxidants that are widely used in stabilization of fish oils as they retard the oxidation of the PUFAs (Hu & Jacobsen, 2016). The antioxidant activity of tocopherols is mainly due to their ability to donate hydrogens to lipid free-radicals (Kamal-Eldin & Appelqvist, 1996). The tocopherol content of the fresh fish oil was: α-tocopherol, 136 ± 0.2 μg/g oil; β-tocopherol, 4 ± 0.3 μg/g oil; γ-tocopherol, 84 ± 0.3 μg/g oil; δ-tocopherol, 37 ± 0.3 μg/g oil. The evolution of tocopherol content during storage is shown in Error! Reference source not found.**B-D**. α-, γ- and δ-tocopherol content at day zero was significantly lower than that of the fresh oil for all seven samples. This indicates that oxidation happened somewhere during processing as mentioned in Section 3.5.1., and that the tocopherols were consumed to inhibit this oxidation. A significant decrease in α-, γ- and δ-tocopherol content between day zero and day 28 was observed for all samples except for δ-tocopherol in the PPH sample. This could indicate that PPH may furthermore contain peptides with antioxidative properties, as previously reported for potato peptides and hydrolysates (Olsen et al., 2020; Yesiltas et al., 2022). Nevertheless, results indicates that none of the samples are completely protected from oxidation as tocopherols are being consumed to prevent the formation of hydroperoxides and secondary oxidation products. In line with the PV results, the tocopherol contents in Pat and PPE at day 28 were significantly lower than the rest of the samples. Decreases of 100, 76-82 and 50% were observed for Pat and PPE after 28 days for α-, γ- and δ-tocopherol, respectively (**Table S11** in Supplementary Material). In comparison, the decrease in α-tocopherol for G1, G38, and PPH was 15, 37 and 30%, respectively. A significant decrease in α-tocopherol from day 0 to day 7 was observed for Pat, PPE, PPH, and Cas. This can be explained, in part, by the lack of a lag-phase in terms of hydroperoxide formation for Pat and PPE.

γ- and δ-tocopherol are primarily being consumed from day 21 to day 28 for Pat and PPE. This could be attributed to the fact that α-tocopherol could have been depleted shortly after day 21 coupled with the large formation of hydroperoxides for Pat and PPE between day 21 and day 28. The difference in the consumption of tocopherols has been attributed to the chemical structures of the molecules. The relative antioxidant activity order of tocopherols in vivo has been reported as α > β > γ > δ (Kamal-Eldin & Appelqvist, 1996). This agrees with the findings of this study.

#### 3.5.3. Secondary oxidation products – volatiles

Hydroperoxides formed during oxidation decompose into secondary volatile oxidation products such as alcohols, ketones, and aldehydes. Due to the presence of double bonds in PUFAs, many different compounds can be formed. These volatile compounds are often considered off-flavors and decrease product quality. Fishy flavor/odor in milk enriched with fish oil has been attributed to a combination of 1-penten-3-one, 4-heptenal, 1-octen-3-one, 1,5-octadien-3-one, 2,4-heptadienal and 2,6-nonadienal (Venkateshwarlu et al., 2004a, 2004b). Three of these compounds ((*Z*)-4-heptenal, 1-penten-3-one and (*E,E*)-2,4-heptadienal) were identified in the microencapsulated fish oil after 28 days of storage. For this study, the evolutions of 1-penten-3-ol, (*E*)-2-pentenal, heptanal, (*Z*)-4-heptenal, (*E,E*)-2,4-heptadienal and octanal during storage were highlighted as these compounds either had high final concentrations or presented different trends. All six compounds have been identified in fish oil (Horiuchi et al., 1998; Jacobsen, 2015). The course of the remaining eight volatiles identified (1-penten-3-one, 2-ethyl-furan, pentanal, 2,3-pentadione, 1-pentanol, 2-hexenal, benzaldehyde, and hexanal) are shown in **Figure S1** in the supplementary materials. The volatiles have odors, which are unacceptable for consumers. These have been identified as fishy and sweet or rancid for (*Z*)-4-heptenal and (*E,E*)-2,4-heptadienal, respectively and chemical or burnt and oily or soapy for heptanal and 2-pentenal, respectively (Hartvigsen et al., 2000). The nasal odor thresholds of the volatiles in oil vary from 0.002 ppm for 4-heptenal to 2.3, 3.2 and 10 ppm for 2-pentenal, heptanal, and 2,4-heptadienal, respectively (Belitz et al., 2009). **Figure 3** shows the development volatiles during storage. In line with the PV and tocopherol results, significantly higher concentrations of (*E*)-2-pentenal, 1-penten-3-ol, heptanal and (*E,E*)-2,4-heptadienal were observed for Pat and PPE compared to the remaining samples. For the same four volatile compounds, there were no significant differences between G1, G38, Cas, PPH and PI after 28 days. This indicates that G1, G38, and PPH are, at least, equal to Cas in terms of oxidative stability. G1 and PPH had lower final concentrations of all six volatiles when compared to Cas while only 1-penten-3-ol and (*Z*)-4-heptenal concentrations were higher for G38 than for Cas. The concentrations of (*E*)-2-pentenal, heptanal and (*E,E*)-2,4-heptadienal were all well below the odor thresholds of the volatiles for G1, G38, and PPH. The lag times in formation of volatiles for G1, G38, PPH and Cas were generally similar, but depended on the compounds. The lag times observed varied from no lag time for octanal to between 2-3 weeks for heptanal. However, the concentration of octanal for G38 at day 14 was not significantly different from day zero, suggesting a lag time of one week. Longer lag times were expected for G1, G38 and PPH compared to the proteins due to their longer PV lag times. The development of octanal, (*E,E*)-2,4-heptadienal and (*Z*)-4-heptenal differs from the other volatile compounds. The octanal formation of Pat and Cas was not significantly different and PI had a significantly lower final concentration of octanal compared to both Cas and G1. The development of volatiles for PI is interesting as it had a significantly higher PV compared to the G1, G38 and PPH while showing the same formation of some of the volatiles ((*E*)-2-pentenal, 1-penten-3-ol and heptanal). The formation of heptanal and (*Z*)-4-heptenal was lowest overall for PI. This shows that even though PI had a higher PV, the formation of these two compounds was still comparable to both the peptides and the control. However, PI had a higher final concentration of both (*E,E*)-2,4-heptadienal and (*E*)-2-pentenal than peptides and the control. (Hermund et al., 2019) and (García-Moreno et al., 2018) both studied the formation of 1-penten-3-ol, (*E*)-2-pentenal and heptanal in fish oil-loaded electrosprayed capsules during 21 days of storage at 20°C. The two studies used different shell materials and a combination of seaweed antioxidants, commercial antioxidants and without antioxidants. (Hermund et al., 2019) reported values above approximately 1000, 350 and 200 ng g^-1^ oil after 21 days of storage for 1-penten-3-ol, (*E*)-2-pentenal and heptanal, respectively. (García-Moreno et al., 2018) reported values above 600, 500 and 200 ng g^-1^ oil for the same compounds. In comparison, the highest concentration of 1-penten-3-ol observed after 21 days in the present study was 266.8 ± 52.9 ng g^-1^ oil for Pat while G1, G38, and PPH were all below 100 ng g^-1^ oil after 28 days. The concentration of (*E*)-2-pentenal and heptanal was below 50 ng g^-1^ for G1, G38, and PPH after 28 days of storage. This shows that the emulsifying peptides and PPH compare favorably to other potential solutions for microencapsulation of fish oil in terms of volatile formation. However, these results must be interpreted with care as the external standards were injected directly onto the Tenax tubes. This means that the results do not consider the potential effect of the food matrix on the release of the volatile compounds.

**Figure 3.**
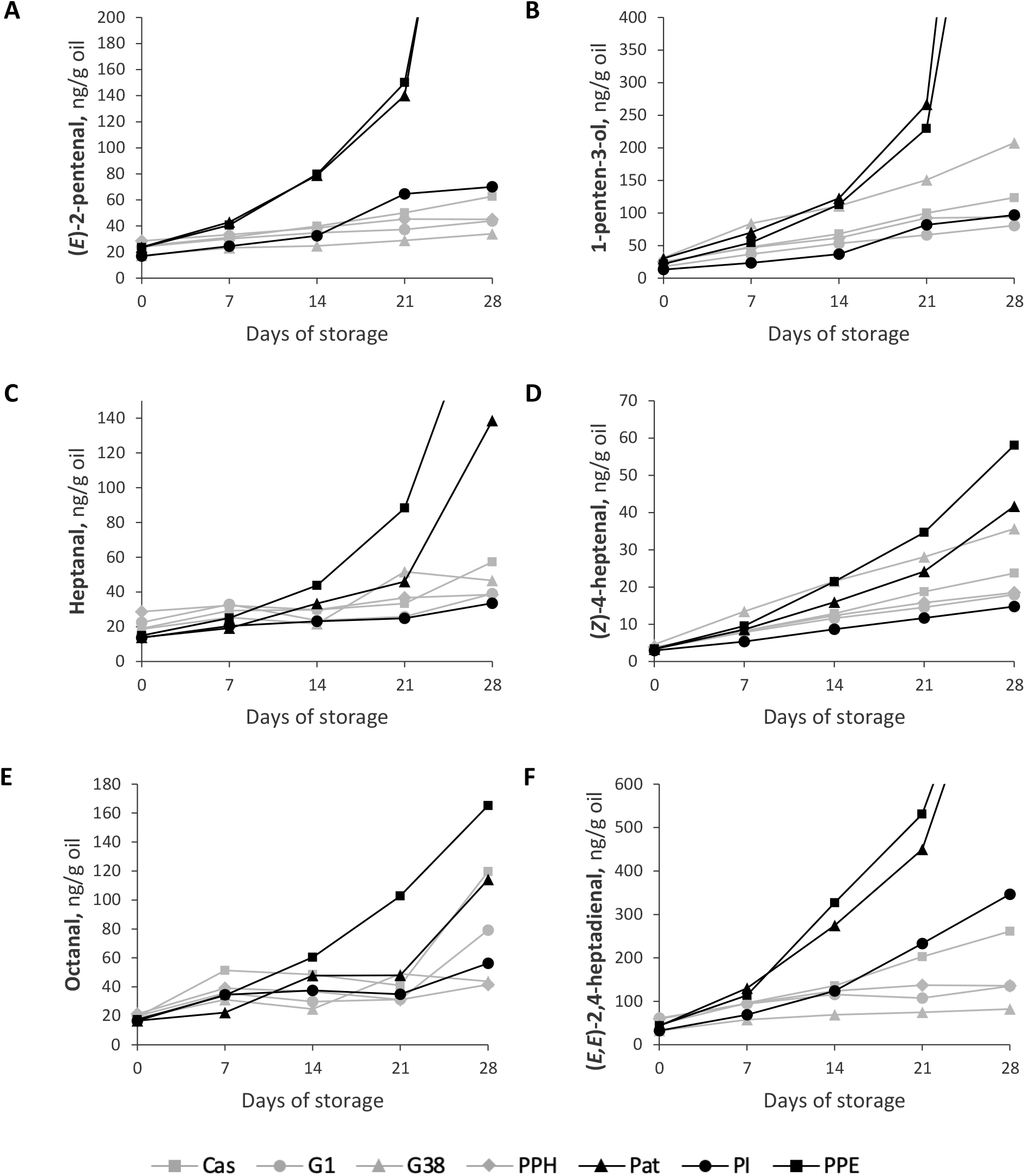
Formation of secondary volatile oxidation products during 28 days of storage for fish-oil, which has been microencapsulated using sodium caseinate, peptides or proteins as emulsifying agents. A) 2-pentenal, B) 1-penten-3-ol, C) Heptanal, D) 4-heptenal, E) Octanal, F) 2,4-heptadienal. Values at day 28 for Pat and PPE can be found in Tables S1 to S6 in the supplementary materials.

There are different plausible explanations as to why peptides and PPH performed better than the other samples in terms of oxidative stability. Proteins can act as antioxidants in different ways (e.g., free radical scavenging or metal chelation). The most reactive amino acids in terms of free radical scavenging are Cys, Met, Phe, Tyr, Trp and His (Elias et al., 2008). G1 contains a single Phe while G38 contains Phe, His and the highly oxidizable Cys. However, as reported by (García-Moreno, Jacobsen, et al., 2020), G1 did not exhibit *in vitro* antioxidative properties (DPPH radical scavenging) when stabilizing fish oil-in-water emulsions compared to G38. They performed similarly in terms of oxidative stability in the present study, thus, different patterns in oxidative stability are observed for emulsions and microencapsulated oil with the same emulsifiers. This indicates that the emulsifying properties of peptides predominantly determines oxidative stability when microencapsulating fish oil. Additionally, previous studies found a positive correlation between the amount of non-encapsulated oil and the formation of primary oxidation products (Velasco et al., 2000). This is in agreement with our findings, as G1, G38, and PPH, which had the highest EEs and performed better in terms of oxidative stability. However, the EE of PI was significantly lower than that of Pat even though the final PV for PI was lower than that of Pat. Other studies conclude that the course of lipid oxidation cannot solely be explained by the amount of non-encapsulated oil. (Drusch & Berg, 2008) investigated the effect of inlet/outlet temperature and oil load on the amount of non-encapsulated oil and the formation of primary and secondary oxidation products. They concluded that the surface oil protects the other fractions of extractable oil (e.g. droplets close to the surface) from oxidation. The non-encapsulated oil may also contain droplets that are surrounded by the peptides which could potentially protect them from oxidation (Drusch et al., 2007). Finally, the presence of free volume in the shell material increases the oxygen diffusivity, which can lead to an increased rate of autoxidation. Free volume is mainly dependent on the molecular weight of the encapsulating agent (e.g., a lower molecular weight results in less free volume) due to a tighter packing of the molecules (Hogan et al., 2001a). Air inclusion during emulsion preparation can also lead to the formation of air bubbles inside the shell material, as observed for PI (**Figure 1F**).

## 4 CONCLUSIONS

The potential of potato proteins and peptides to be used as emulsifiers in microencapsulation of fish oil was studied. Emulsions stabilized with the synthetic peptides G1 and G38 and the potato protein hydrolysate had lower oil droplet sizes and exhibited better physical stability compared to emulsions stabilized with intact/parent potato proteins. Spherical microcapsules with minimal agglomeration were produced for both proteins and peptides, however, with varying capsule diameters and amounts of non-encapsulated oil. Encapsulation efficiencies of microcapsules obtained with G1, G38, and PPH were higher than those produced with sodium caseinate and the proteins, indicating that even at a low concentration of 0.2 wt%, the peptides and hydrolysate show high potential and performs better than sodium caseinate at the same concentration. The oxidative stability followed the trends in encapsulation efficiency and the microcapsules obtained with peptides and hydrolysate showed better oxidative stability compared to those produced with their parent proteins. The differences in oxidative stability may be attributed to several factors, but the ability of the proteins and peptides to form physically stable emulsions being the most important. In all aspects of the study, the emulsifying peptides performed as good as, or better than sodium caseinate, indicating that potato peptides can be used as efficient emulsifiers in microencapsulation of hydrophobic bioactive ingredients. Interestingly, the hydrolysate performed similarly to the individual peptides in all aspects. This work shows that a valuable functional ingredient can be obtained by utilizing targeted hydrolysis of potato protein extract, even without the need for further processing.

## 5 Acknowledgements

This work was part of Protein valorization through informatics, hydrolysis, and separation (PROVIDE) project, which is supported by Innovation Fund Denmark (grant number: 7045-00021B). The authors would also like to thank Lihme Protein Solutions (Kgs. Lyngby, Denmark) and Dennis Hansen for providing the purified extracts of patatin and protease inhibitors. Finally, we want to tank KMC AmbA (Brande, Denmark) for providing samples.

## SUPPLEMENTARY MATERIALS

**Table S1.** 1-penten-3-ol

|  | 0 | 7 | 14 | 21 | 28 |
| --- | --- | --- | --- | --- | --- |
| Caseinate | 25 ± 3.0<sup>c,v</sup> | 48 ± 6.0<sup>ab,vw</sup> | 68 ± 10<sup>b,w</sup> | 100 ± 10<sup>ab,x</sup> | 124 ± 16<sup>a,x</sup> |
| G1 | 18 ± 2.2<sup>ab,v</sup> | 37 ± 2.6<sup>a,w</sup> | 53 ± 4.3<sup>ab,x</sup> | 66 ± 6.9<sup>a,y</sup> | 81 ± 4.7<sup>a,z</sup> |
| G38 | 31 ± 1.0<sup>d,v</sup> | 84 ± 21<sup>c,w</sup> | 111 ± 15<sup>c,w</sup> | 151 ± 9.1<sup>b,x</sup> | 208 ± 12<sup>a,y</sup> |
| PPH | 25 ± 0.3<sup>c,v</sup> | 47 ± 8.0<sup>ab,w</sup> | 62 ± 0.9<sup>ab,w</sup> | 92 ± 4.8<sup>ab,x</sup> | 94 ± 11<sup>a,x</sup> |
| Patatin | 30 ± 1.0<sup>d,v</sup> | 79 ± 5.3<sup>bc,vw</sup> | 122 ± 17<sup>c,w</sup> | 267 ± 53<sup>c,x</sup> | 1424 ± 142<sup>c,z</sup> |
| PI | 13 ± 0.8<sup>a,v</sup> | 19 ± 7.8<sup>a,vw</sup> | 37 ± 3.7<sup>a,w</sup> | 82 ± 8.0<sup>a,x</sup> | 97 ± 4.5<sup>a,y</sup> |
| PPE | 22 ± 2.9<sup>bc,v</sup> | 55 ± 10<sup>ab,w</sup> | 113 ± 5.6<sup>c,x</sup> | 230 ± 13<sup>c,y</sup> | 1007 ± 31<sup>b,z</sup> |

Letters (a-e) indicate statistically significant differences ( $p < 0.05$ ) between samples.
Letters (v-z) indicate statistically significant differences ( $p < 0.05$ ) between the days.

**Table S2.** Heptanal

|  | 0 | 7 | 14 | 21 | 28 |
| --- | --- | --- | --- | --- | --- |
| Caseinate | 19 ± 0.9<sup>b,v</sup> | 29 ± 0.4<sup>bc,vw</sup> | 30 ± 2.2<sup>b,vw</sup> | 33 ± 4.6<sup>b,w</sup> | 57 ± 8.8<sup>a,x</sup> |
| G1 | 23 ± 0.3<sup>c,v</sup> | 33 ± 4.3<sup>c,w</sup> | 24 ± 1.1<sup>a,v</sup> | 26 ± 1.1<sup>a,v</sup> | 39 ± 2.3<sup>a,w</sup> |
| G38 | 19 ± 1.1<sup>b,v</sup> | 25 ± 2.7<sup>ab,v</sup> | 22 ± 0.4<sup>a,v</sup> | 52 ± 0.9<sup>c,w</sup> | 47 ± 14<sup>a,w</sup> |
| PPH | 29 ± 1.7<sup>d,v</sup> | 32 ± 2.4<sup>c,vw</sup> | 30 ± 0.4<sup>b,v</sup> | 37 ± 1.4<sup>b,vw</sup> | 39 ± 6.1<sup>a,w</sup> |
| Patatin | 14 ± 1.3<sup>a,v</sup> | 19 ± 0.1<sup>a,w</sup> | 33 ± 1.3<sup>b,x</sup> | 46 ± 1.8<sup>c,y</sup> | 138 ± 8.9<sup>b,z</sup> |
| PI | 14 ± 1.4<sup>a,v</sup> | 16 ± 7.8<sup>a,vw</sup> | 23 ± 0.7<sup>a,w</sup> | 25 ± 0.4<sup>a,w</sup> | 34 ± 4.5<sup>a,x</sup> |
| PPE | 15 ± 1.6<sup>a,v</sup> | 25 ± 1.4<sup>ab,w</sup> | 44 ± 2.3<sup>c,x</sup> | 88 ± 1.2<sup>d,y</sup> | 226 ± 12<sup>c,z</sup> |

Letters (a-e) indicate statistically significant differences ( $p < 0.05$ ) between samples.
Letters (v-z) indicate statistically significant differences ( $p < 0.05$ ) between the days.

**Table S3.**
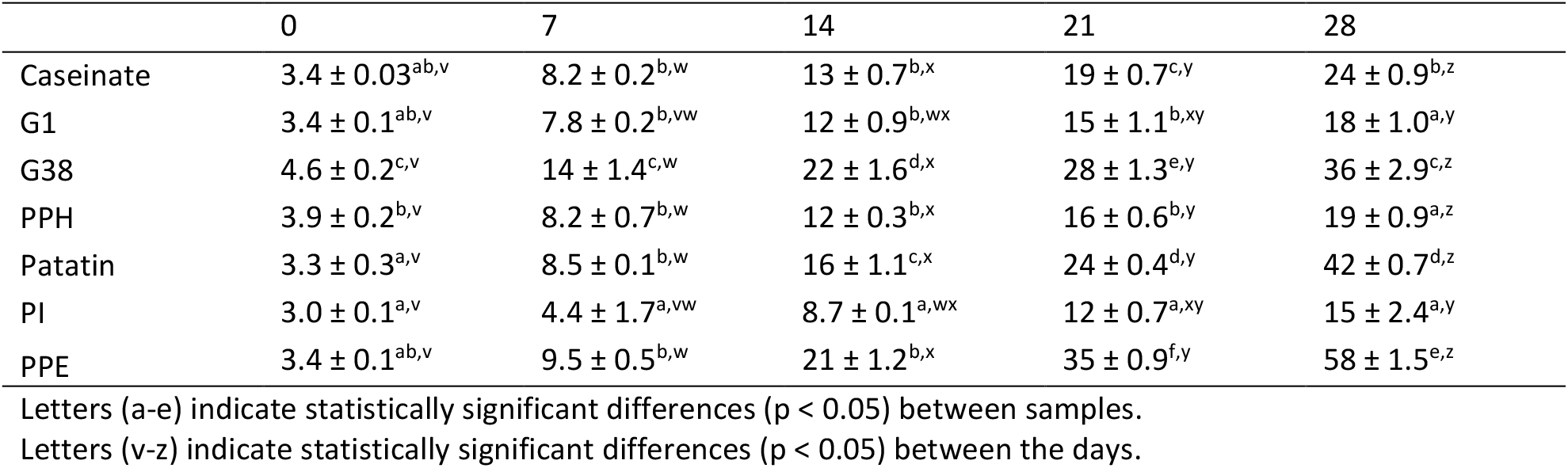
4-heptenal

|  | 0 | 7 | 14 | 21 | 28 |
| --- | --- | --- | --- | --- | --- |
| Caseinate | 3.4 ± 0.03<sup>ab,v</sup> | 8.2 ± 0.2<sup>b,w</sup> | 13 ± 0.7<sup>b,x</sup> | 19 ± 0.7<sup>c,y</sup> | 24 ± 0.9<sup>b,z</sup> |
| G1 | 3.4 ± 0.1<sup>ab,v</sup> | 7.8 ± 0.2<sup>b,vw</sup> | 12 ± 0.9<sup>b,wx</sup> | 15 ± 1.1<sup>b,xy</sup> | 18 ± 1.0<sup>a,y</sup> |
| G38 | 4.6 ± 0.2<sup>c,v</sup> | 14 ± 1.4<sup>c,w</sup> | 22 ± 1.6<sup>d,x</sup> | 28 ± 1.3<sup>e,y</sup> | 36 ± 2.9<sup>c,z</sup> |
| PPH | 3.9 ± 0.2<sup>b,v</sup> | 8.2 ± 0.7<sup>b,w</sup> | 12 ± 0.3<sup>b,x</sup> | 16 ± 0.6<sup>b,y</sup> | 19 ± 0.9<sup>a,z</sup> |
| Patatin | 3.3 ± 0.3<sup>a,v</sup> | 8.5 ± 0.1<sup>b,w</sup> | 16 ± 1.1<sup>c,x</sup> | 24 ± 0.4<sup>d,y</sup> | 42 ± 0.7<sup>d,z</sup> |
| PI | 3.0 ± 0.1<sup>a,v</sup> | 4.4 ± 1.7<sup>a,vw</sup> | 8.7 ± 0.1<sup>a,wx</sup> | 12 ± 0.7<sup>a,xy</sup> | 15 ± 2.4<sup>a,y</sup> |
| PPE | 3.4 ± 0.1<sup>ab,v</sup> | 9.5 ± 0.5<sup>b,w</sup> | 21 ± 1.2<sup>b,x</sup> | 35 ± 0.9<sup>f,y</sup> | 58 ± 1.5<sup>e,z</sup> |

Letters (a-e) indicate statistically significant differences ( $p < 0.05$ ) between samples.
Letters (v-z) indicate statistically significant differences ( $p < 0.05$ ) between the days.

**Table S4.** 2,4-heptadienal

|  | 0 | 7 | 14 | 21 | 28 |
| --- | --- | --- | --- | --- | --- |
| Caseinate | 47 ± 4.0<sup>abc,v</sup> | 96 ± 1.0<sup>bc,vw</sup> | 136 ± 21<sup>b,w</sup> | 202 ± 16<sup>bc,x</sup> | 261 ± 32<sup>b,y</sup> |
| G1 | 61 ± 2.9<sup>c,v</sup> | 95 ± 8.0<sup>bc,vw</sup> | 116 ± 11<sup>ab,vw</sup> | 108 ± 45<sup>a,vw</sup> | 135 ± 21<sup>a,w</sup> |
| G38 | 32 ± 3.8<sup>a,v</sup> | 58 ± 6.2<sup>a,vw</sup> | 69 ± 4.8<sup>a,w</sup> | 75 ± 7.9<sup>a,w</sup> | 82 ± 18<sup>a,w</sup> |
| PPH | 59 ± 10<sup>bc,v</sup> | 95 ± 18<sup>bc,vw</sup> | 124 ± 13<sup>ab,w</sup> | 137 ± 17<sup>ab,w</sup> | 136 ± 33<sup>a,w</sup> |
| Patatin | 43 ± 7.5<sup>ab,v</sup> | 130 ± 2.5<sup>d,w</sup> | 274 ± 37<sup>c,x</sup> | 450 ± 48<sup>d,y</sup> | 1039 ± 4.5<sup>c</sup> |
| PI | 32 ± 4.1<sup>a,v</sup> | 53 ± 28<sup>ab,vw</sup> | 124 ± 7.1<sup>ab,w</sup> | 233 ± 15<sup>c,x</sup> | 346 ± 59<sup>b,y</sup> |
| PPE | 44 ± 6.9<sup>abc,v</sup> | 113 ± 20<sup>cd,v</sup> | 327 ± 35<sup>c,w</sup> | 531 ± 37<sup>d,x</sup> | 1081 ± 19<sup>c,z</sup> |

Letters (a-e) indicate statistically significant differences ( $p < 0.05$ ) between samples.
Letters (v-z) indicate statistically significant differences ( $p < 0.05$ ) between the days.

**Table S5.**
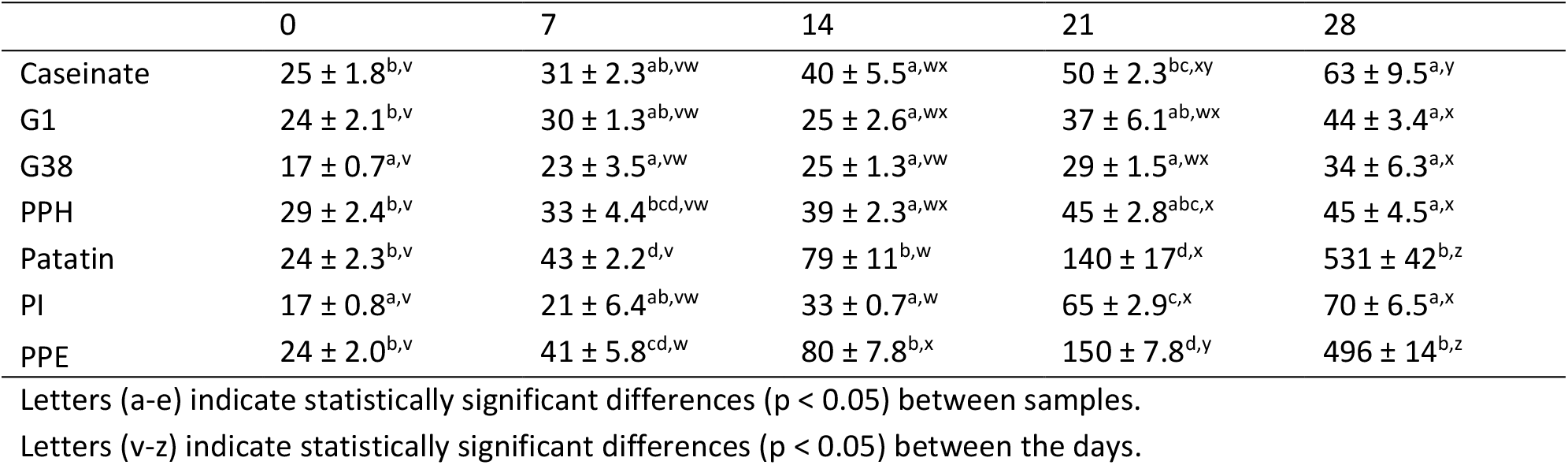
2-Pentenal

**Table S6.** Octanal

|  | 0 | 7 | 14 | 21 | 28 |
| --- | --- | --- | --- | --- | --- |
| Caseinate | 19 ± 0.8<sup>bcd,v</sup> | 51 ± 0.8<sup>e,x</sup> | 49 ± 3.7<sup>c,wx</sup> | 41 ± 1.3<sup>b,w</sup> | 120 ± 5.2<sup>c,y</sup> |
| G1 | 20 ± 0.3<sup>cd,v</sup> | 36 ± 1.5<sup>cd,w</sup> | 30 ± 1.3<sup>a,w</sup> | 31 ± 2.3<sup>a,w</sup> | 79 ± 4.4<sup>b,x</sup> |
| G38 | 19 ± 1.0<sup>abc,v</sup> | 31 ± 1.7<sup>b,w</sup> | 25 ± 0.2<sup>a,v</sup> | 49 ± 0.1<sup>c,x</sup> | 44 ± 4.9<sup>a,x</sup> |
| PPH | 21 ± 1.0<sup>d,v</sup> | 39 ± 2.8<sup>d,x</sup> | 37 ± 0.7<sup>b,x</sup> | 31 ± 2.0<sup>a,w</sup> | 42 ± 2.6<sup>a,x</sup> |
| Patatin | 17 ± 1.5<sup>a,v</sup> | 22 ± 0.8<sup>a,w</sup> | 48 ± 2.5<sup>c,x</sup> | 48 ± 0.6<sup>c,x</sup> | 114 ± 6.1<sup>c,z</sup> |
| PI | 17 ± 0.1<sup>a,v</sup> | 28 ± 12<sup>bcd,w</sup> | 38 ± 1.1<sup>b,w</sup> | 35 ± 0.7<sup>a,w</sup> | 56 ± 8.6<sup>a,x</sup> |
| PPE | 17 ± 0.2<sup>ab,v</sup> | 34 ± 1.0<sup>bc,w</sup> | 60 ± 3.5<sup>d,x</sup> | 103 ± 2.5<sup>d,y</sup> | 165 ± 6.8<sup>d,z</sup> |

Letters (a-e) indicate statistically significant differences ( $p < 0.05$ ) between samples.
Letters (v-z) indicate statistically significant differences ( $p < 0.05$ ) between the days.

**Table S7.** Alpha-tocopherol

|  | 0 | 7 | 14 | 21 | 28 |
| --- | --- | --- | --- | --- | --- |
| Caseinate | 20 ± 0.5<sup>cd,x</sup> | 13 ± 2.4<sup>ab,v</sup> | 18 ± 0.0<sup>a,wx</sup> | 21 ± 0.7<sup>d,x</sup> | 15 ± 0.4<sup>d,vw</sup> |
| G1 | 21 ± 1.1<sup>cd,v</sup> | 18 ± 0.5<sup>b,v</sup> | 20 ± 3.0<sup>a,v</sup> | 21 ± 0.1<sup>d,v</sup> | 18 ± 0.1<sup>e,v</sup> |
| G38 | 21 ± 0.4<sup>cd,v</sup> | 16 ± 0.2<sup>b,v</sup> | 19 ± 3.5<sup>a,v</sup> | 17 ± 0.0<sup>c,v</sup> | 13 ± 0.4<sup>c,v</sup> |
| PPH | 23 ± 1.0<sup>d,y</sup> | 17 ± 0.5<sup>b,vw</sup> | 18 ± 0.0<sup>a,wx</sup> | 20 ± 0.2<sup>d,x</sup> | 16 ± 0.2<sup>d,v</sup> |
| Patatin | 18 ± 0.1<sup>ab,y</sup> | 14 ± 0.7<sup>ab,x</sup> | 13 ± 1.2<sup>a,x</sup> | 8.3 ± 1.3<sup>b,w</sup> | ND<sup>a,v</sup> |
| PI | 20 ± 0.1<sup>bc,v</sup> | 16 ± 3.2<sup>b,v</sup> | 14 ± 6.7<sup>a,v</sup> | 16 ± 0.2<sup>c,v</sup> | 10 ± 0.1<sup>b,v</sup> |
| PPE | 16 ± 0.1<sup>a,y</sup> | 9.4 ± 0.0<sup>a,x</sup> | 9.4 ± 0.8<sup>a,x</sup> | 3.7 ± 0.6<sup>a,w</sup> | ND<sup>a,v</sup> |

Letters (a-e) indicate statistically significant differences ( $p < 0.05$ ) between samples.
Letters (v-z) indicate statistically significant differences ( $p < 0.05$ ) between the days.

**Table S8.** Gamma-tocopherol

|  | 0 | 7 | 14 | 21 | 28 |
| --- | --- | --- | --- | --- | --- |
| Caseinate | 14 ± 0.7<sup>ab,w</sup> | 12 ± 0.4<sup>a,v</sup> | 10.9 ± 0.0<sup>b,v</sup> | 14.6 ± 0.1<sup>e,w</sup> | 11.5 ± 0.1<sup>b,v</sup> |
| G1 | 14 ± 0.1<sup>ab,w</sup> | 12 ± 0.3<sup>a,v</sup> | 10.7 ± 0.1<sup>b,v</sup> | 14.2 ± 0.2<sup>de,w</sup> | 12.1 ± 0.2<sup>b,v</sup> |
| G38 | 14 ± 0.3<sup>ab,x</sup> | 12 ± 0.1<sup>a,w</sup> | 10.3 ± 0.2<sup>b,v</sup> | 13.0 ± 0.0<sup>c,wx</sup> | 10.4 ± 0.2<sup>b,v</sup> |
| PPH | 15 ± 0.9<sup>b,w</sup> | 12 ± 0.3<sup>a,v</sup> | 10.6 ± 0.1<sup>b,v</sup> | 14.4 ± 0.2<sup>e,w</sup> | 11.8 ± 0.1<sup>b,v</sup> |
| Patatin | 13 ± 0.1<sup>a,w</sup> | 11 ± 0.4<sup>a,w</sup> | 10.7 ± 0.5<sup>b,w</sup> | 11.6 ± 0.2<sup>b,w</sup> | 2.1 ± 1.4<sup>a,v</sup> |
| PI | 13 ± 0.3<sup>a,v</sup> | 11 ± 2.4<sup>a,v</sup> | 10.4 ± 0.8<sup>ab,v</sup> | 13.7 ± 0.0<sup>cd,v</sup> | 10.1 ± 0.2<sup>b,v</sup> |
| PPE | 12 ± 0.2<sup>a,x</sup> | 9.2 ± 0.3<sup>a,w</sup> | 9.2 ± 0.2<sup>a,w</sup> | 9.3 ± 0.0<sup>a,w</sup> | 3.0 ± 0.3<sup>a,v</sup> |

Letters (a-e) indicate statistically significant differences ( $p < 0.05$ ) between samples.
Letters (v-z) indicate statistically significant differences ( $p < 0.05$ ) between the days.

**Table S9.** Delta-tocopherol

|  | 0 | 7 | 14 | 21 | 28 |
| --- | --- | --- | --- | --- | --- |
| Caseinate | 5.7 ± 0.1<sup>ab,v</sup> | 5.4 ± 0.5<sup>a,v</sup> | 5.1 ± 0.2<sup>a,v</sup> | 5.9 ± 0.6<sup>a,v</sup> | 5.0 ± 0.0<sup>b,v</sup> |
| G1 | 5.8 ± 0.1<sup>ab,v</sup> | 5.1 ± 0.1<sup>a,v</sup> | 5.0 ± 0.0<sup>a,v</sup> | 5.2 ± 0.6<sup>a,v</sup> | 5.0 ± 0.1<sup>b,v</sup> |
| G38 | 6.0 ± 0.0<sup>b,y</sup> | 5.7 ± 0.2<sup>a,x</sup> | 4.9 ± 0.0<sup>a,wx</sup> | 5.0 ± 0.0<sup>a,vw</sup> | 4.5 ± 0.1<sup>b,v</sup> |
| PPH | 5.7 ± 0.5<sup>ab,v</sup> | 5.1 ± 0.3<sup>a,v</sup> | 4.9 ± 0.2<sup>a,v</sup> | 5.4 ± 0.2<sup>a,v</sup> | 5.1 ± 1.1<sup>b,v</sup> |
| Patatin | 5.5 ± 0.2<sup>a,w</sup> | 4.9 ± 0.2<sup>a,w</sup> | 5.0 ± 0.4<sup>a,w</sup> | 4.6 ± 0.1<sup>a,w</sup> | 2.7 ± 0.2<sup>a,v</sup> |
| PI | 5.5 ± 0.0<sup>ab,v</sup> | 4.8 ± 1.1<sup>a,v</sup> | 5.0 ± 0.1<sup>a,v</sup> | 5.3 ± 0.2<sup>a,v</sup> | 4.5 ± 0.0<sup>b,v</sup> |
| PPE | 5.6 ± 0.0<sup>ab,x</sup> | 4.8 ± 0.1<sup>a,w</sup> | 4.6 ± 0.1<sup>a,w</sup> | 5.0 ± 0.2<sup>a,w</sup> | 2.8 ± 0.1<sup>a,v</sup> |

Letters (a-e) indicate statistically significant differences ( $p < 0.05$ ) between samples.
Letters (v-z) indicate statistically significant differences ( $p < 0.05$ ) between the days.

**Table S10.** Peroxide value

|  | 0 | 7 | 14 | 21 | 28 |
| --- | --- | --- | --- | --- | --- |
| Caseinate | 1.5 ± 0.0<sup>ab,v</sup> | 3.6 ± 0.8<sup>a,vw</sup> | 6.0 ± 1.1<sup>ab,wx</sup> | 8.3 ± 0.6<sup>a,x</sup> | 15 ± 1.5<sup>a,y</sup> |
| G1 | 2.0 ± 0.1<sup>abc,v</sup> | 3.4 ± 0.1<sup>a,vw</sup> | 7.7 ± 2.2<sup>ab,x</sup> | 4.4 ± 0.5<sup>a,vwx</sup> | 6.0 ± 0.2<sup>a,wx</sup> |
| G38 | 1.1 ± 0.3<sup>a,v</sup> | 3.7 ± 0.5<sup>a,v</sup> | 4.1 ± 0.3<sup>a,vw</sup> | 5.0 ± 0.3<sup>a,vw</sup> | 9.0 ± 2.8<sup>a,w</sup> |
| PPH | 2.6 ± 0.2<sup>c,v</sup> | 3.4 ± 0.1<sup>a,vw</sup> | 5.0 ± 0.2<sup>a,wx</sup> | 6.4 ± 0.6<sup>a,x</sup> | 8.9 ± 0.6<sup>a,y</sup> |
| Patatin | 2.4 ± 0.2<sup>bc,v</sup> | 9.4 ± 1.6<sup>b,w</sup> | 19 ± 1.5<sup>c,x</sup> | 32 ± 4.3<sup>c,y</sup> | 188 ± 28<sup>c</sup> |
| PI | 1.5 ± 0.6<sup>ab,v</sup> | 4.3 ± 0.2<sup>a,w</sup> | 9.3 ± 0.1<sup>b,x</sup> | 22 ± 0.7<sup>b,y</sup> | 33 ± 0.4<sup>a,z</sup> |
| PPE | 2.8 ± 0.1<sup>c,v</sup> | 13 ± 0.5<sup>c,w</sup> | 21 ± 0.4<sup>c,x</sup> | 34 ± 0.7<sup>c,y</sup> | 127 ± 4.4<sup>b</sup> |

Letters (a-e) indicate statistically significant differences ( $p < 0.05$ ) between samples.
Letters (v-z) indicate statistically significant differences ( $p < 0.05$ ) between the days.

**Table S11.** Percentage-wise decrease in tocopherol content after 28 days of storage, %

|  | Alpha-tocopherol | Gamma-tocopherol | Delta-tocopherol |
| --- | --- | --- | --- |
| Caseinate | 24.4 | 16.7 | 10.7 |
| G1 | 14.8 | 12.9 | 13.0 |
| G38 | 36.5 | 23.0 | 24.2 |
| PPH | 30.3 | 22.9 | 10.0 |
| Patatin | 100.0 | 83.2 | 51.3 |
| PI | 48.7 | 24.6 | 18.3 |
| PPE | 100.0 | 75.8 | 50.3 |

**Figure S1.**
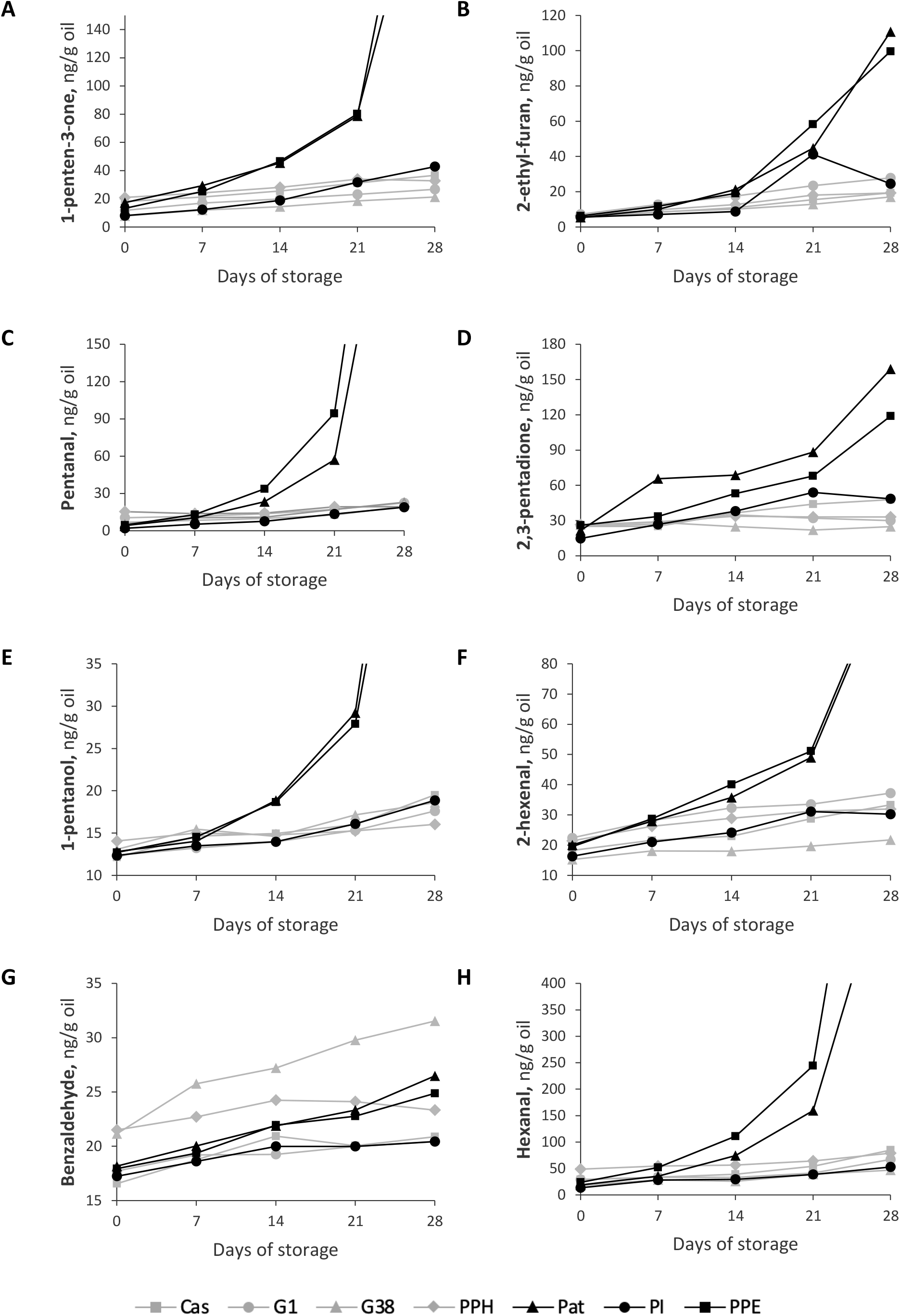
Secondary volatile oxidation products. A) 1-penten-3-one, B) 2-ethyl-furan, C) Pentanal, D) 2,3-pentadione, E) 1-pentanol, F) 2-hexenal, G) Benzaldehyde, H) Hexanal

**Figure S2.**
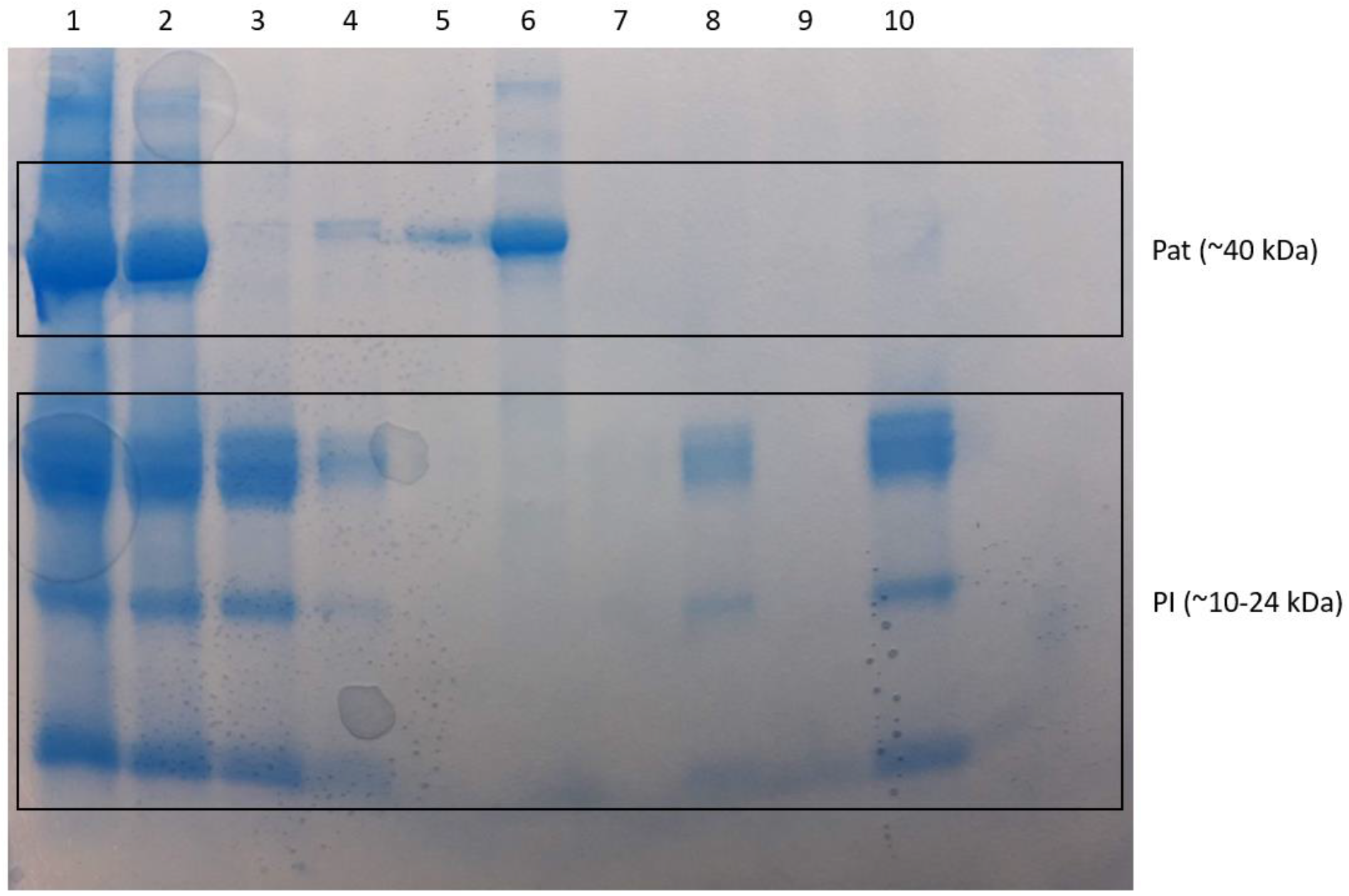
SDS-PAGE analysis of potato protein extract (PPE), patatin-rich fraction (Pat), and protease inhibitor-rich fraction (PI). Well 1 to 6 is a work-up of Pat, showing initial PPE (1), intermediate precipitates (2-5), and the final Pat fraction (6). Wells 7 to 10 is a work-up of PI, showing the intermediary supernatants after NaCl wash (7-9) and the final PI fraction (10). PI was processed from an intermediate precipitate (well 3) and not directly from the PPE.

